# A fast mrMLM algorithm for multi-locus genome-wide association studies

**DOI:** 10.1101/341784

**Authors:** Cox Lwaka Tamba, Yuan-Ming Zhang

## Abstract

**Background:** Recent developments in technology result in the generation of big data. In genome-wide association studies (GWAS), we can get tens of million SNPs that need to be tested for association with a trait of interest. Indeed, this poses a great computational challenge. There is a need for developing fast algorithms in GWAS methodologies. These algorithms must ensure high power in QTN detection, high accuracy in QTN estimation and low false positive rate.

**Results:** Here, we accelerated mrMLM algorithm by using GEMMA idea, matrix transformations and identities. The target functions and derivatives in vector/matrix forms for each marker scanning are transformed into some simple forms that are easy and efficient to evaluate during each optimization step. All potentially associated QTNs with P-values ≤ 0.01 are evaluated in a multi-locus model by LARS algorithm and/or EM-Empirical Bayes. We call the algorithm FASTmrMLM. Numerical simulation studies and real data analysis validated the FASTmrMLM. FASTmrMLM reduces the running time in mrMLM by more than 50%. FASTmrMLM also shows high statistical power in QTN detection, high accuracy in QTN estimation and low false positive rate as compared to GEMMA, FarmCPU and mrMLM. Real data analysis shows that FASTmrMLM was able to detect more previously reported genes than all the other methods: GEMMA/EMMA, FarmCPU and mrMLM.

**Conclusions:** FASTmrMLM is a fast and reliable algorithm in multi-locus GWAS and ensures high statistical power, high accuracy of estimates and low false positive rate.

**Author Summary:** The current developments in technology result in the generation of a vast amount of data. In genome-wide association studies, we can get tens of million markers that need to be tested for association with a trait of interest. Due to the computational challenge faced, we developed a fast algorithm for genome-wide association studies. Our approach is a two stage method. In the first step, we used matrix transformations and identities to quicken the testing of each random marker effect. The target functions and derivatives which are in vector/matrix forms for each marker scanning are transformed into some simple forms that are easy and efficient to evaluate during each optimization step. In the second step, we selected all potentially associated SNPs and evaluated them in a multi-locus model. From simulation studies, our algorithm significantly reduces the computing time. The new method also shows high statistical power in detecting significant markers, high accuracy in marker effect estimation and low false positive rate. We also used the new method to identify relevant genes in real data analysis. We recommend our approach as a fast and reliable method for carrying out a multi-locus genome-wide association study.

## Background

Genome-wide association studies (GWAS) aim at investigating the genetic foundation of complex traits by focusing on the relationship between molecular markers and these traits [1, 2]. Initially, each accession was genotyped by SSR markers, and only hundreds of SSR markers were used to conduct GWAS in plants [3]. When restriction association site DNA sequencing (RAD-seq), specific-locus amplified fragment sequencing (SLAF-seq) and gene chip technology were adopted, then, there were tens of thousands single nucleotide polymorphisms (SNPs) available. When deep sequencing is implemented, millions of SNPs are obtained. As third generation sequencing technology generates, we can get tens of millions SNPs. Evidently, severe computational challenges are faced. Accordingly, there is a critical need for in-depth study of fast algorithm in GWAS methodologies.

Mixed linear model (MLM) method of GWAS was firstly established by Zhang et al. [4]. At each putative QTN scan in this approach, the pedigree-based coancestry matrices of QTN and polygenes are incorporated into the mixed linear model framework to estimate three variance components of QTN, polygenes and residual error. Its long running time makes this method unfashionable. Yu et al. [5] replaced the pedigree-based coancestry matrix with kinship matrix (**K**) to define the degree of genetic covariance between pairs of individuals and view QTN effect as fixed. This method was improved by the spectral decomposition of **H** = **ZKZ**′ + *δ***I** and joint maximum likelihood estimation of fixed effect **β** and 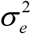, where **Z** is an incidence matrix mapping each observed phenotype to each inbred strain, **I** is an identity matrix and *δ* is the ratio of residual 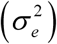 to polygenic 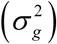 variances. This is efficient mixed model association (EMMA) of Kang et al. [6]. Several approaches have been considered with the aim to reduce the computing time and increase power in QTN detection. Zhang et al. [7] suggested ‘population parameters previously determined’ (P3D) where individuals in polygenic effect are replaced by their corresponding groups, and kinship among individuals is replaced by the kinship among groups. The P3D eliminates the need to re-compute variance components. Kang et al. [8] also fixed the above *δ* in EMMA eXpedited (EMMAX). Li et al. [9] optimized the combination of kinship algorithms and grouping algorithms. Also, other authors suggested alternatives, such as FaST-LMM [10], FaST-LMM-Select [11], GRAMMAR-Gamma [12] and SUPER [13]. In genome-wide EMMA (GEMMA) of Zhou & Stephens [14], especially, target functions and derivatives in vector/matrix forms for each marker, despite their complicated appearance, are easy and efficient to evaluate during each optimization step. Among the above fast methods, the SNP effect was treated as being fixed. Although a random marker model has several advantages [15, 16], an efficient computational algorithm to estimate SNP effect needs to be addressed.

Multi-locus model has become the state-of-the-art GWAS procedure. Previous studies have reported higher power of QTN detection in multi-locus models as compared to the single-marker GWAS analysis [17, 16]. Bonferroni multiple test correction in the single-marker analysis is replaced by a less stringent selection criterion in multi-locus GWAS analysis [16, 18]. Therefore significant loci for complex traits are not missed out. Initially, many statistical approaches were used to estimate all the SNP effects in multi-locus model, such as Bayesian LASSO [19], penalized logistic regression [20, 21], adaptive mixed LASSO [22], elastic net [23], empirical Bayes [24], and empirical Bayes LASSO [25]. If the number of SNPs is large, the above methods are not feasible, even through the Bayesian sparse linear mixed model of Zhou et al. [26] and Bayesian mixture model of Moser et al. [27]. To overcome this shortcoming, Segura et al. [17] proposed multi-locus mixed-model (MLMM), which is a simple, stepwise mixed-model regression with forward inclusion and backward elimination. In FarmCPU of Liu et al. [28], the fixed effect model and random effect model are used iteratively until a stage of convergence is reached. The fixed effect model contains the testing marker and pseudo QTNs to control false positives. The pseudo QTNs are selected from associated markers and evaluated by the random effect model, with k defined by the pseudo-QTNs. However, the computationally intensive forward-backward inclusion of SNPs is clearly a limiting factor in exploring the huge model space. Recently, we have proposed several multi-locus two-stage GWAS approaches, such as mrMLM [16], FASTmrEMMA [18] and ISIS EM-BLASSO [29]. In the first stage, single-locus methods are used to scan all the markers on the genome. In the second stage, a few SNPs potentially associated with the trait are selected and placed into the multi-locus model, and all the effects in the model are estimated by empirical Bayes for true QTN detection. Among our three methods, it is possible to quicken mrMLM.

In this study, we accelerated mrMLM algorithm of Wang et al. [16] using the GEMMA idea and matrix transformation of Miller [30]. In other words, target functions and derivatives in vector/matrix forms for each marker are transformed into some simple forms that are easy and efficient to evaluate during each optimization step. We call this method FASTmrMLM. A series of Monte Carlo simulation experiments and real data analyses were used to validate FASTmrMLM. FASTmrMLM significantly reduces the running time of the mrMLM algorithm. High power and low false positive rate (FPR) in QTN detection and high accuracy in QTN effect estimation were also observed as compared to GEMMA, FarmCPU and mrMLM.

## Results

### Computational efficiency

To confirm the effectiveness of FASTmrMLM, a series of Monte Carlo simulation experiments were carried out. Each sample was analyzed by five methods: FASTmrMLM, mrMLM, FarmCPU, GEMMA, and EMMA. In the first Monte Carlo simulation experiment where no polygenic variance was simulated, the running times (Intel Core i5-4570 CPU 3.20GHz, Memory 7.88G) for the above five methods are 6.25, 13.77, 5.12, 2.57 and 68.77 (hours), respectively (Fig 1 and Table 1). FASTmrMLM takes less than 50% of the running time needed when mrMLM is used. The same trend is observed in the rest of the simulations (Fig 1 and Table 1). It indicates that FASTmrMLM significantly quickens mrMLM. Although GEMMA and FarmCPU have lower computational time than FASTmrMLM, their performances in statistical power and parameter estimation accuracy are worse than those of FASTmrMLM.

**Fig 1.**
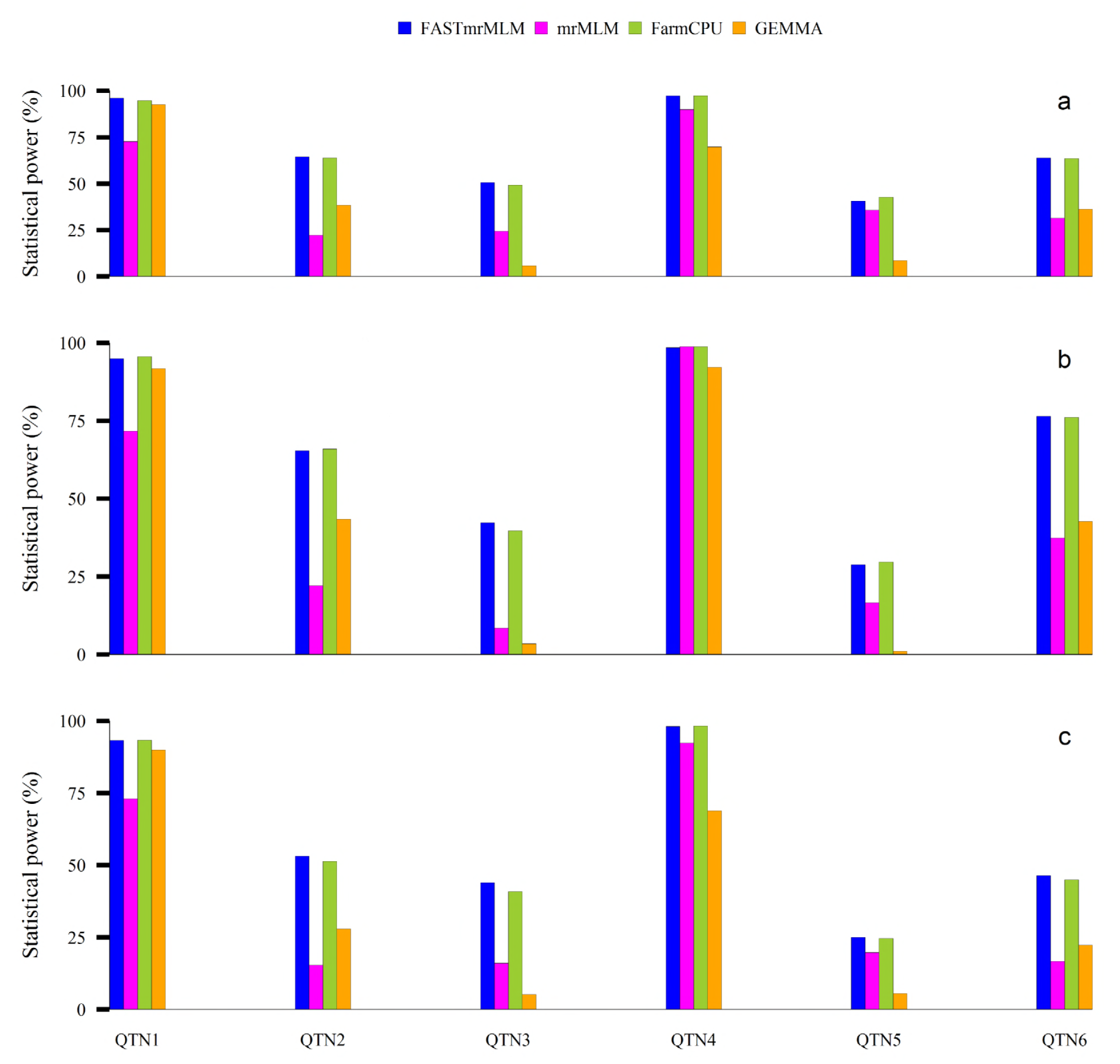
Computing times (hours) taken by FASTmrMLM, mrMLM, FarmCPU, GEMMA, and EMMA in the Monte Carlo simulation experiments I (**a**), II (**b**) and III (**c**).

**Table 1.**
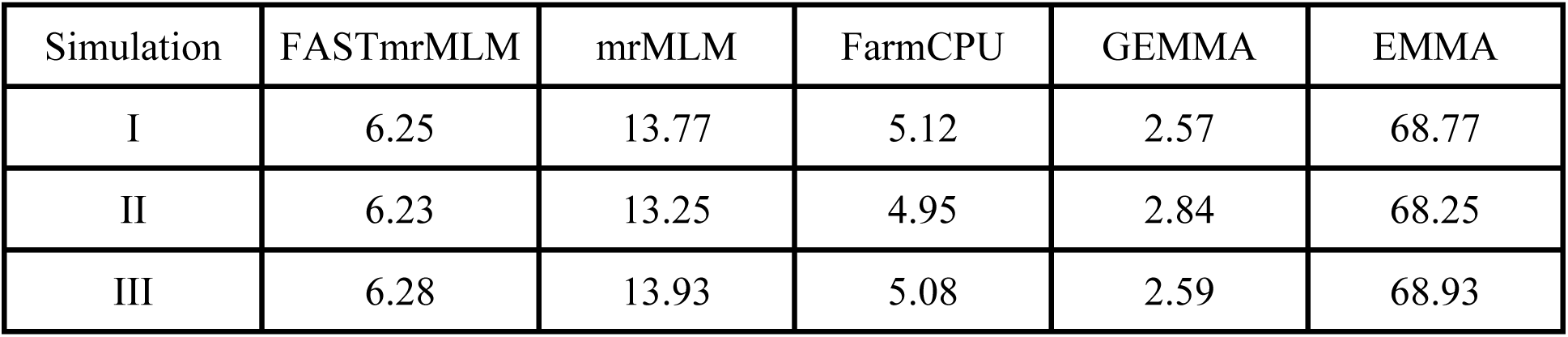
Comparison of the time taken (Hrs) in the detection of QTNs in three simulation experiments using FASTmrMLM, GEMMA/EMMA, FarmCPU and mrMLM methods (199 individuals each with 10000 SNPs, 1000 replicates)

### Statistical power

Statistical power was used to evaluate the effectiveness of FASTmrMLM when compared with the other four methods. All the statistical powers in the three simulation experiments are presented in Fig 2 and Table S1. In the first simulation experiment (Fig 2a and Table S1), the average powers across six simulated QTN from FASTmrMLM, mrMLM, FarmCPU, GEMMA and EMMA were 68.8, 68.6, 41.9, 46.0 and 46.0 (%), respectively. It means that FASTmrMLM has the highest power in QTN detection. When paired t-test was conducted between FASTmrMLM and the other methods, FASTmrMLM has significantly higher power than FarmCPU, GEMMA, and EMMA (P-value = 0.004 ~ 0.012). Although there is no significant difference between FASTmrMLM and mrMLM (P-value = 0.688), FASTmrMLM has slightly higher power than mrMLM (Table S2). Therefore, FASTmrMLM is the most effective method for QTN detection.

**Fig 2.**
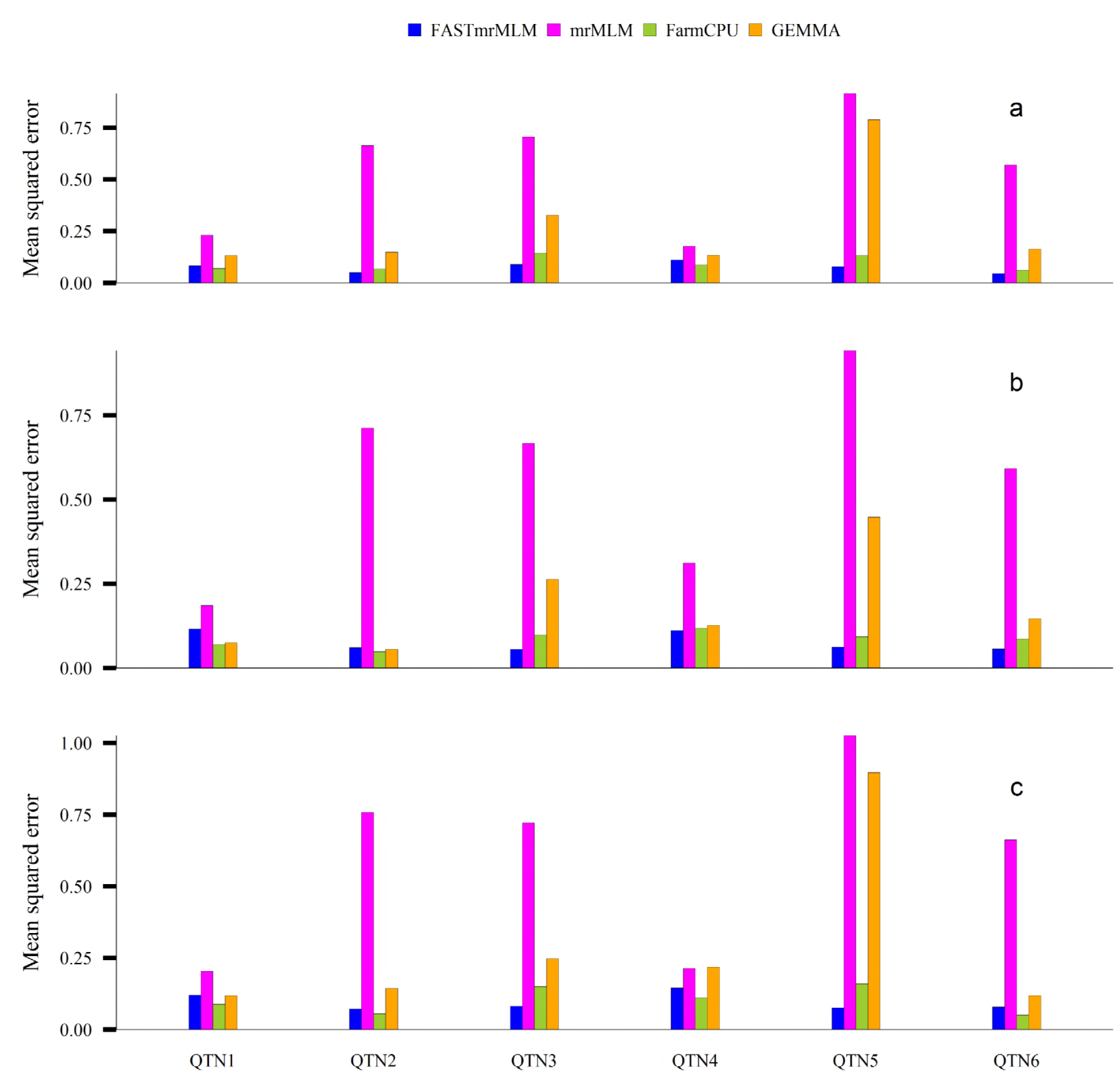
Statistical powers for all the simulated QTNs detected by FASTmrMLM, mrMLM, FarmCPU, and GEMMA in Monte Carlo simulation experiments I (**a**), II (**b**) and III (**c**)

### Mean squared errors of estimated QTN effects

Mean squared error (MSE) was used to measure the accuracy of each estimated QTN effect for all the five methods in the three simulation experiments. All the MSE values for all the six simulated QTN effects in all the three simulation experiments are shown in Fig 3 and Table S3, and the average value for each simulated QTN effect in all the three simulation experiments is listed in Table S4. In the first simulation experiment (Fig 3a and Table S3), average MSE values across six simulated QTN from FASTmrMLM, mrMLM, FarmCPU, GEMMA, and EMMA were 0.0775, 0.0933, 0.2824, 0.5467 and 0.5432, respectively. It means that FASTmrMLM has the highest accuracy in the estimation of QTN effect. When paired t-test was conducted between FASTmrMLM and the other four methods, the MSE value was at least significantly lower from FASTmrMLM than from GEMMA and EMMA (P-value = 0.009 ~ 0.020). Although there is no significant difference between FASTmrMLM and the other two (mrMLM and FarmCPU) methods (P-value = 0.110 ~ 0.806), FASTmrMLM has slightly lower MSE than mrMLM and FarmCPU (Table S2). Therefore, FASTmrMLM has the highest accuracy in the estimation of QTN effect.

**Fig 3.**
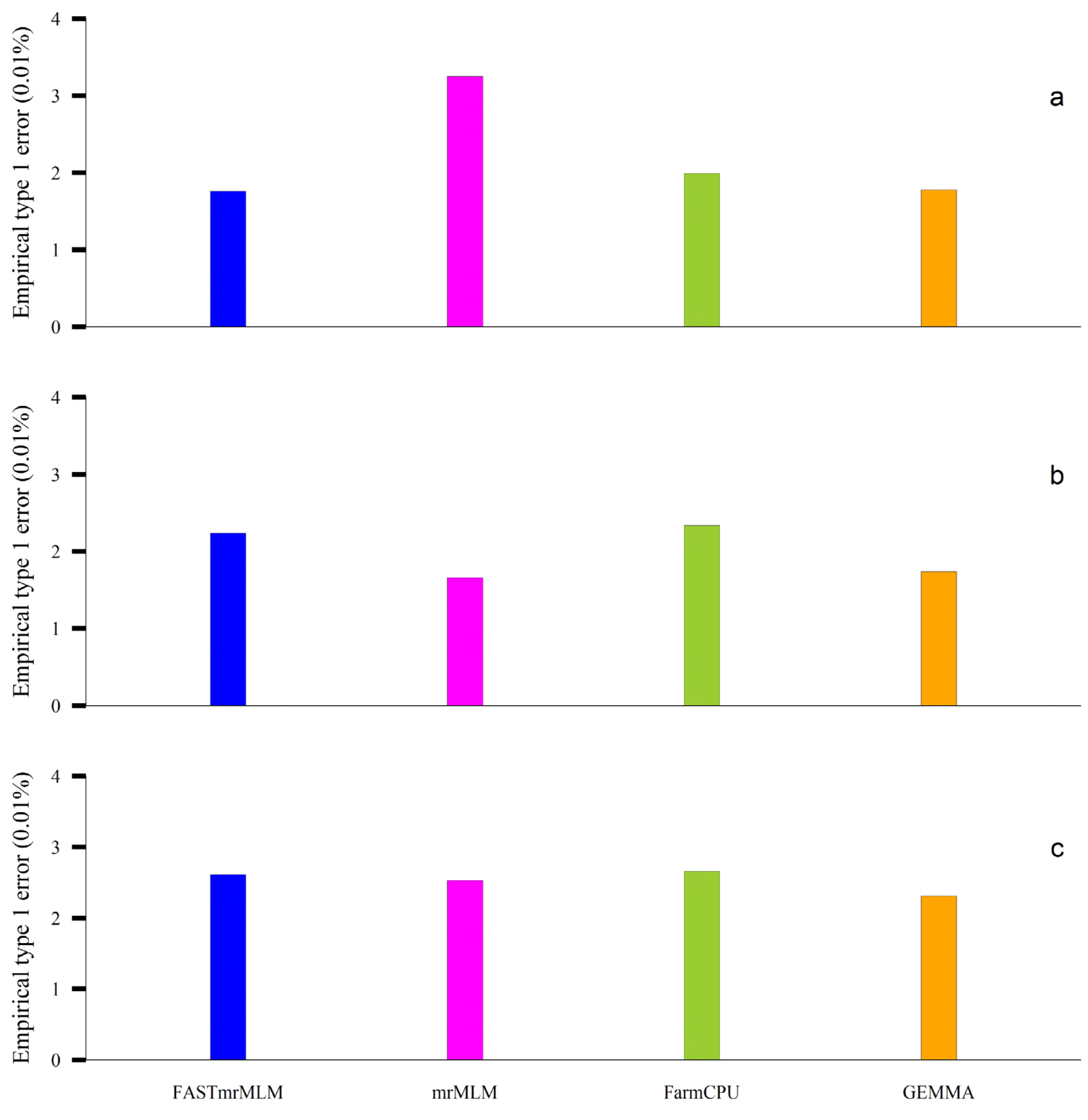
Mean square error (MSE) for all the simulated QTN effects estimated by F FASTmrMLM, mrMLM, FarmCPU, and GEMMA in Monte Carlo simulation experiments I (**a**), II (**b**) and III (**c**).

### False positive rate

All the single-locus GWAS approaches are involved in multiple test issue so that Bonferroni correction is frequently used to control false positive rate (FPR) or empirical type 1 errors. In the current multi-locus GWAS method, a less stringent selection criterion was used to identify true QTN. In this situation, it is important to show the FPR values in the three simulation experiments. All the FPR values are presented in Fig 4 and Table 2. In the first simulation experiment (Fig 4a and Table 2), the FPR values for the above five methods were 1.80E-2, 1.99E-2, 1.78E-2, 3.25E-2 and 3.25E-2 (%), respectively (Table 2). It indicates that FASTmrMLM has almost the lowest FPR, although a less stringent selection criterion is adopted. Therefore, FASTmrMLM has controlled the FPR in QTN detection.

**Fig 4.**
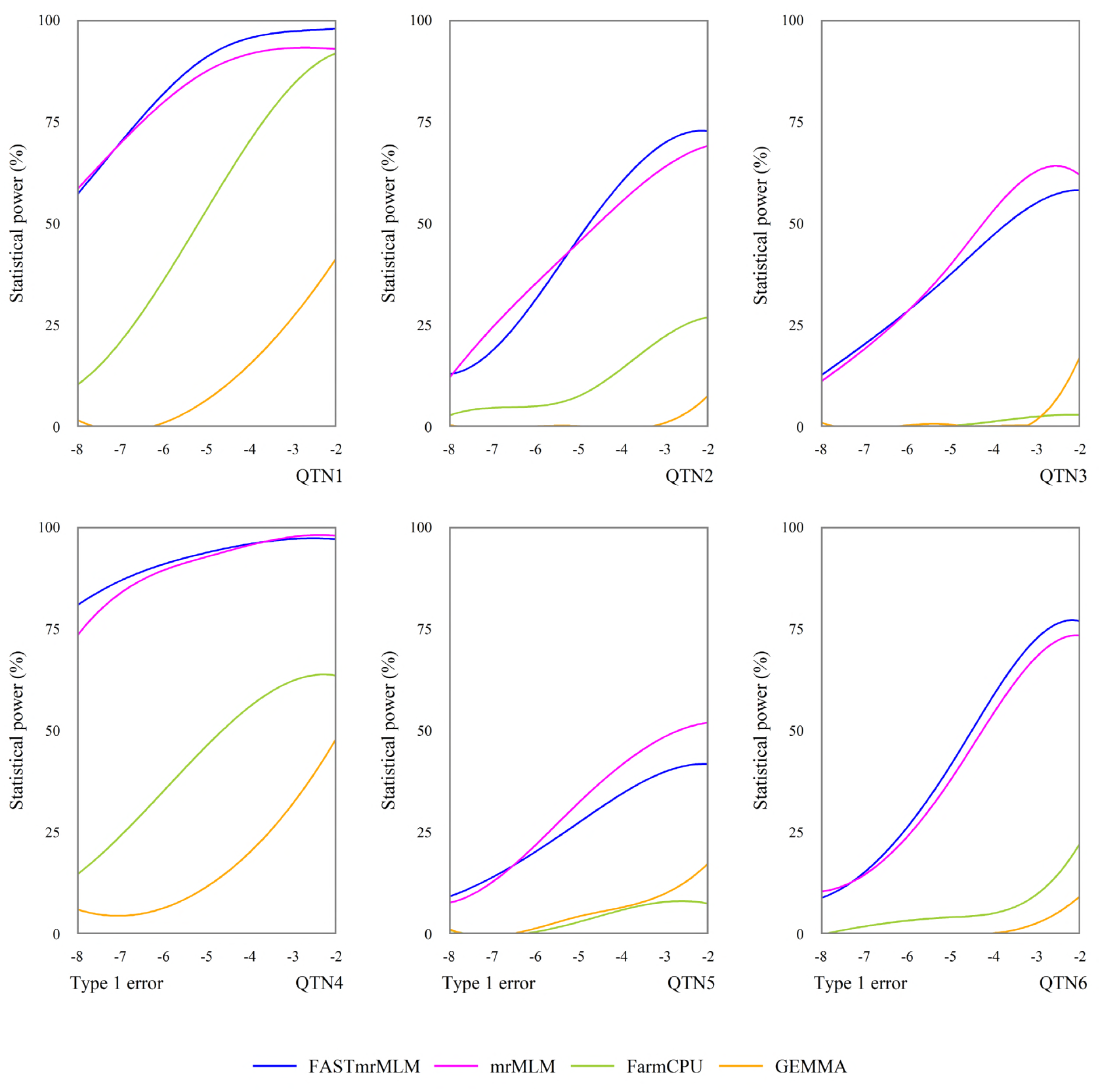
Empirical type 1 error rates derived from FASTmrMLM, mrMLM, FarmCPU, and GEMMA in the Monte Carlo simulation experiments I (**a**), II (**b**) and III (**c**).

**Table 2.** Comparison of false positive rate (%) in the detection of QTNs in three simulation experiments using FASTmrMLM, mrMLM, FarmCPU, and GEMMA/EMMA methods

| Simulation | FASTmrMLM | mrMLM | FarmCPU | GEMMA | EMMA |
| --- | --- | --- | --- | --- | --- |
| I | 0.1799 | 0.1990 | 0.1780 | 0.3250 | 0.3250 |
| II | 0.2276 | 0.2340 | 0.1740 | 0.1660 | 0.1660 |
| III | 0.2611 | 0.2660 | 0.2310 | 0.2530 | 0.2530 |

### Receiver operating characteristic curve

Receiver operating characteristic (ROC) curve is obtained when statistical power is plotted against controlled Type 1 error. This curve is used to compare the efficiencies of different methods in the detection of significant effects. A method is considered the best if its ROC curve lies above all the other curves of the methods being compared. We simulated 100 various probability levels of significance between 1E-8 to 1E-2. We calculated statistical powers corresponding to these levels in the first simulation experiment. Fig 5 depicts the ROC curves for the four methods for each simulated QTN in the first experiment. As a result, FASTmrMLM performs the best for the simulated QTNs 1, 2, 4 and 6; mrMLM has the highest curve for QTNs 3 and 5 though its curves lie slightly above ROC curves of FASTmrMLM.

**Fig 5.**
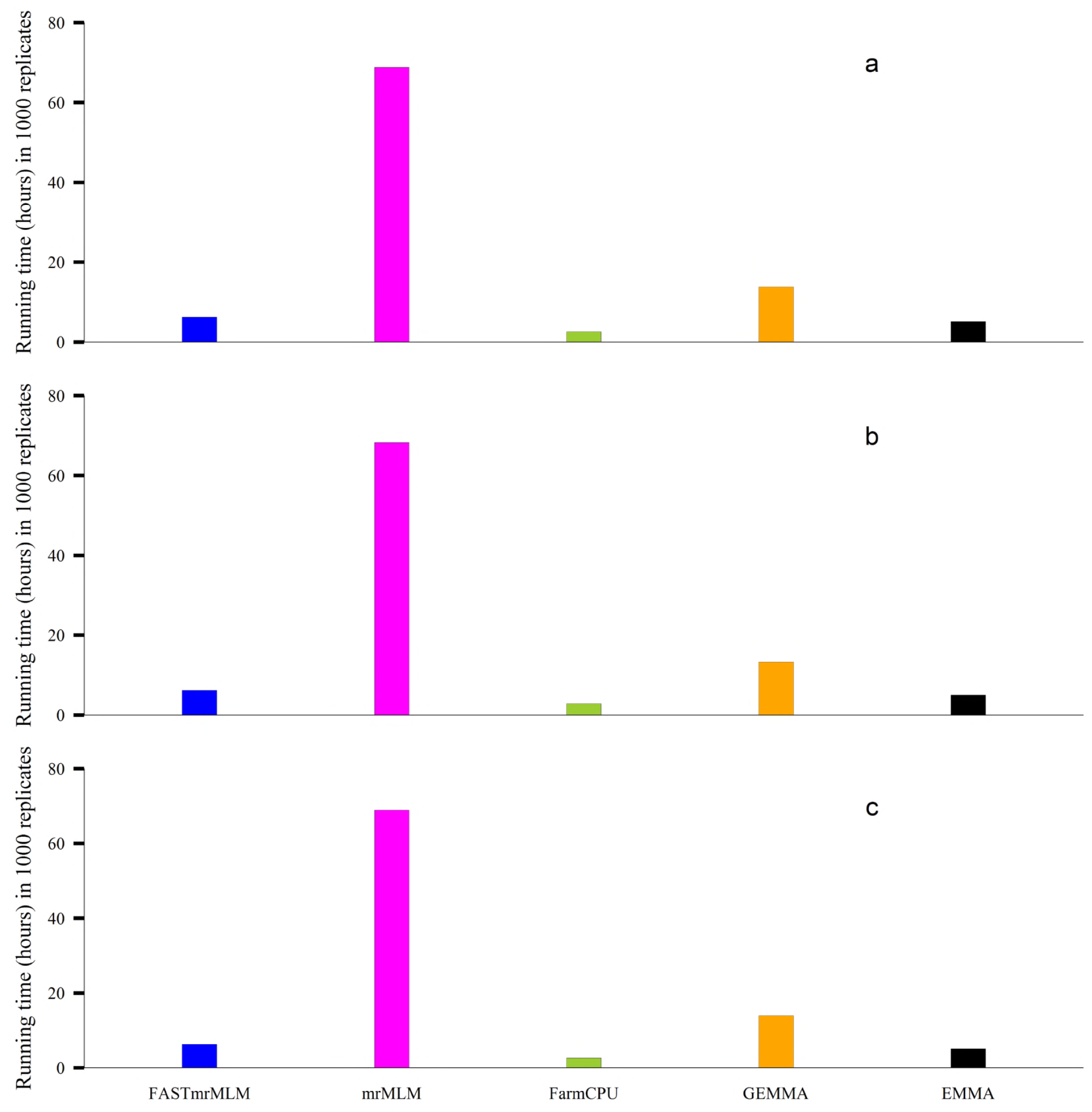
Statistical powers of all the simulated QTNs in the first simulation experiment plotted against Type 1 error (in a log10 scale) for the five GWAS methods (FASTmrMLM, mrMLM, FarmCPU, and GEMMA).

### Real data analysis

We used FASTMRMLM, mrMLM, FarmCPU, and GEMMA/EMMA methods to re-analyze six Arabidopsis flowering time traits (LD, LDV, SD, 0W, 2W, and 4W) in Atwell et al. [31]. FASTmrMLM identified 17, 15, 14, 17, 14 and 15 SNPs to be significantly associated respectively with the six traits above. The identified SNPs for each trait were used to conduct a multiple linear regression analysis, and we calculated the corresponding AIC and BIC values. Table S5 shows the AIC and BIC values for all the methods in all the six traits. FASTmrMLM has low AIC and BIC values for nearly all the traits. mrMLM compares almost equally with FASTmrMLM. This indicates that SNPs found to be significant by FASTmrMLM fit the data better than the other methods.

We obtained 14, 10, 7, 10, 10, and 11 known genes in the proximity of detected SNPs when we used FASTmrMLM method to analyze the six traits respectively. The mrMLM method identified 4, 5, 1, 2, 5, and 4 known genes respectively for the above six traits. Known genes identified by FarmCPU for the six traits were 2, 3, 2, 0, 0, 4 and 2 respectively. GEMMA/EMMA was able to determine 5, 2, 0, 0, 3, and 3 known genes for the above six traits. FASTmrMLM detected more known genes than all the other methods (Table S6).

The new method was able to detect 7, 7, 6, 8, 7 and 5 new genes for the corresponding six traits considered in this study (Table S7). GEMMA/EMMA was not able to detect any gene for the traits SD and 0W. Indeed the Bonferroni correction is so stringent and may make significant genes to be missed it. The same observations are made when FarmCPU is used to analyze the 0W trait, and no gene is identified. From the results obtained in this study, we observe that the new approach can detect more associated genes when used in GWAS study. FASTmrMLM does reduce the computational time as well as ensures that the associated genes are not missed out. Based on these findings, we note that *Arabidopsis thaliana* GWAS results obtained in this study are reliable.

## Discussion

FASTmrMLM in this study is different from mrMLM of Wang et al. [16] in two aspects. First, computations during each optimization stage in the first stage of FASTmrMLM are quicker than those in mrMLM. This is because FASTmrMLM uses simplified analytic forms of gradient functions and Hessian matrices, and mrMLM uses numerical gradients and Hessian matrices during optimization stage. The simplified analytic forms in target functions and derivatives quicken computation during each optimization stage of FASTmrMLM. Our results confirm the significant reduction in running time in FASTmrEMMA. Second, the LARS algorithm of Efron et al. [32] is used in FASTmrMLM to select the *n* − 1 variables that are most likely associated with the quantitative trait of interest if the number of markers passing the 1% level of significance is more than *n* in the second stage of FASTmrMLM. The *n* − 1 selected markers are then included in a multi-locus model and detected by EM-Empirical Bayes for true QTN identification. This can explain why slight improvements in statistical power and accuracy are observed in this study. In this study we also confirmed why we used EM-Empirical Bayes to estimate QTN effects in the multi-locus model. Here we compared three methods: EM-Empirical Bayes, adaptive Lasso, and SCAD. As a result, EM-Empirical Bayes has higher statistical power and more accurate than SCAD and adaptive Lasso (Tables S8 to S10).

In the past ten years, many approaches have been used to reduce running time in GWAS. First, QTN effect is viewed as fixed [5, 7, 8] rather than as previously viewed as random [4]. As such, the number of variance components decreases from three (QTN, polygenic and residual error) to two (polygenic and residual error). Second, the polygenic-to-residual variance ratio, obtained at null hypothesis, is fixed in the single-marker genome scan [7, 8]. In this case, the number of variance components decreases further from two (polygenic and residual error) to one (residual error). Finally, some matrix transformations and identities are adopted. One such matrix transformation is spectral decomposition, which lets target functions and derivatives be expressed by simple forms, such as in Kang et al. [6] and Wen et al. [18]. The first stage of the new method (FASTmrMLM) considers a QTN effect as random. Then, the polygenic-to-residual variance ratio obtained at null hypothesis is fixed in the single-marker genome testing. With a simple matrix transformation, the matrix h turns to be a sum of two matrices (an identity matrix (**I**) and a matrix of rank one (**Z**_*ci*_**Z**_*ci*_’*ω_i_*)). More importantly, the results in Miller [30] are then used to compute quickly **H**^−1^, **|H|** and quadratic terms in the form **η′H**^−1^**τ**, which frequently appear in the target functions, derivatives and parameter optimization in Newton-Raphson iterations.

In the single-locus scanning step of FASTmrMLM, all estimates are based on restricted maximum likelihood estimation (REML). This is because FASTmrMLM is an extension of mrMLM, which only considers REML estimates [16]. Of course, we preferred REML over maximum likelihood estimation because it produces unbiased estimates of the variance components by taking into account the degrees of freedom that result from evaluating the fixed effects.

As showed in this study, FASTmrMLM is better than GEMMA. The possible reasons are described below. First, FASTmrMLM considers QTN effect as random rather than as fixed as in GEMMA of Zhou & Stephens [14]. This confirms the advantages of a random marker model as outlined in Goddard et al. [15]. Then, the significance level for each test is LOD=3 in FASTmrMLM rather than 0.05/*m* in GEMMA. In theory, a less stringent selection criterion simultaneously increases statistical power and FPR. However, our new method increases not only statistical power but also FPR. Finally, FASTmrMLM is a multi-locus model method while GEMMA is a single-locus model approach. This can explain why various significance levels are adopted between FASTmrMLM and GEMMA.

## Conclusion

We accelerated our previous multi-locus GWAS method: mrMLM, with the help of a new matrix transformation and matrix identities. As a result, the computational time of estimating variance components in the first step is significantly reduced. We implemented the LARS algorithm of Efron et al. [32] between the first step and EM-Empirical Bayes estimation in the second step. This makes slight improvements in statistical power and estimation accuracy as compared to mrMLM. We confirmed that EM-Empirical Bayes is the best method for the estimation of parameters in the multi-locus model. The proposed method, named FASTmrMLM, significantly increased the statistical power and decreased FPR compared with other methods: FarmCPU, GEMMA and EMMA. In real data analysis, more previously reported genes were detected by FASTmrMLM.

## Materials and Methods

### Genetic model

We consider a mixed linear regression model,

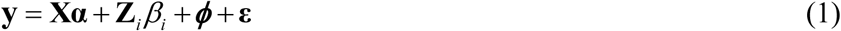

with **y** being a *n*×1 phenotypic vector of quantitative trait of *n* individuals, **X** is an *n* × *q* incident matrix of fixed effects **α** including the overall mean, **Z**_i_ is an *n* ×1 vector of the zth SNP, *β_i_* is a random effect of the *i*th marker, it is assumed to be a normal distribution with zero mean and each marker prior variance 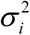, 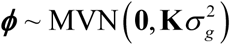 is the polygenic effect with a multivariate normal distribution with zero mean and variance 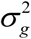 described by a covariance matrix **K** (kinship matrix), and 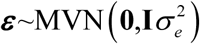 is the residual error **I** with being *n*×*n* identity matrix. In this study, the kinship matrix **K** is marker inferred kinship [33] defined as 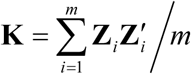.

From equation (1) we have that,

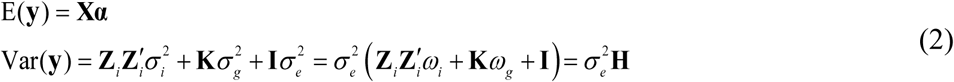

where 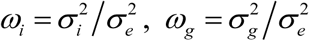 are the variance ratios and 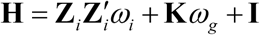. In this model, we have three random effects: the *i*th marker effect, polygenic effect, and residual error. Under the pure polygenic model, we have that E(**y**) = **X*α*** and 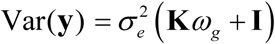.

We pre-estimate the value of 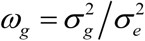 under the pure polygenic model and fix it when testing each SNP effect in the genome-wide scanning [7]. Using spectral decomposition, we can find a diagonal matrix **D** = diag(*δ*_1_,.L, *δ_m_*) and a matrix **U** such that **K** = **UDU**′. Notice that spectral decomposition can be performed on **K** since as defined it is a square, symmetric matrix. Transforming **y** in Equation (1) by multiplying by **U**′, we have,

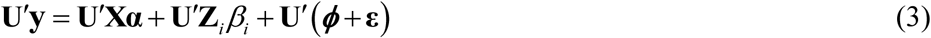

Let 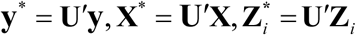, then Equation (3) becomes

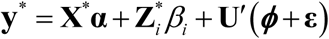

It follows that,

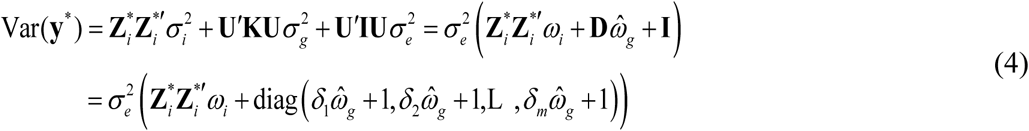

For simplicity let 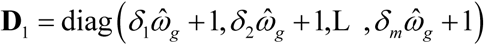. 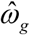 is fixed and **D**_1_ as defined is a positive semi-definite matrix. Therefore, we can obtain 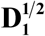. Further transforming **y**\* by multiplying equation (3) through by 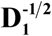 and letting 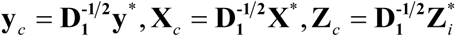 and 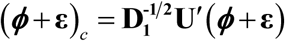. We have that,

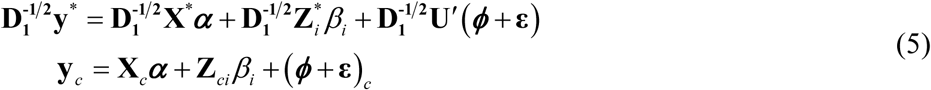

Notice that the transformation in Equation (5) is equivalent to multiplying the original **y** by 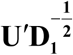 i.e. 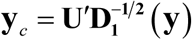. Now,

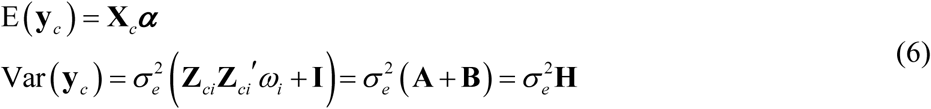

with **A** = **Z**_*ci*_**Z***_ci_′ω_i_*, **B** = **I**, and **H** = **A** + **B**. Thus the distribution of our transformed data vector **y**_*c*_ is normal with the mean **X**_*c*_**α** and variance-covariance matrix, 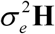. The parameters to be estimated 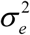 are *ω_i_* and for *i* = 1, 2, L, m.

### Parameter estimation

The profiled residual log likelihood (REML) of **y**\* after absorbing other terms in the constant term *C* is,

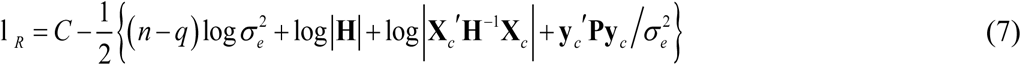

where, **P** = **H**^−1^ − **H**^−1^**X**_*c*_ (**X_*c*_**′**H**^−1^**X**_*c*_)^−1^ **X**_*c*_^′^**H**^−1^, and *q =* rank (**X**_*c*_). We differentiate Equation (7) to obtain REML estimates as shown below.

**Estimation of residual variance** 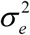: We have

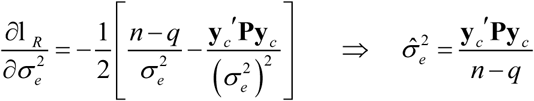

This can be simplified to

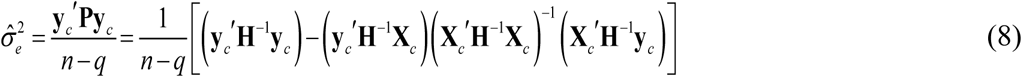

**Estimation of the variance ratio** *ω_i_*: We have that,

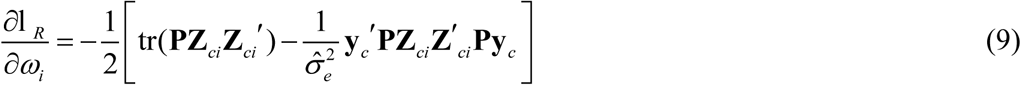

The second derivative of the residual log likelihood function with respect to is given below, and it is used to obtain the Hessian matrix.

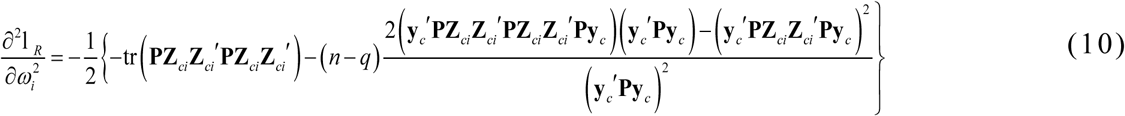

Evaluation of REML equation and its derivatives require **H**^−1^ and |**H|**. **H** is a sum of two matrices. The inverse and determinant of **B** can easily be computed because it is an identity matrix, and **A** is a matrix of rank 1. In mrMLM of Wang et al. [16], **H** is also a sum of two matrices but in this new algorithm (FASTmrMLM) we have simplified the **H** further to be the sum of an identity matrix and a matrix of rank one. Therefore,

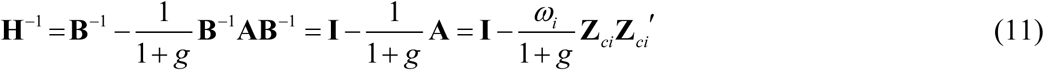

and, |**H**| = (1 + *g*) |**B**| = 1 + *g*, where *g =* tr(**AB**^−1^) = tr(**A**) = *ω_i_* tr (**Z**_*ci*_**Z**_*ci*_′) [30]. REML requires many quadratic terms in the form **η**′**H**^−1^**τ**, which can be expressed as

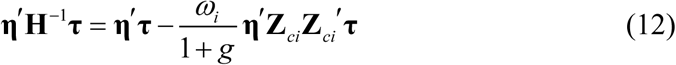

where **η**′ and **τ** can be any vectors or matrices with **H**^−1^ such as **X**_c_′**H**^−1^**X**_*c*_, **y**_*c*_′**Py**_*c*_ or **X**_*c*_′**Py**_*c*_. Note that 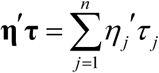, where *η_j_* corresponds to the *j*th row (element) of the matrix (vector) **η** and *τ_j_* corresponds to the *j*th row (element) of the matrix (vector) **τ** for *j* = 1,2,L, n. Notice that the value in Equation (8) above can easily be estimated because with the help of Equation (12) each term in the bracket in Equation (8) is in the form **η**′**H**^−1^**τ**. For example, 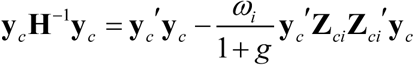. Also, the gradient function/score function in Equation (9) and Hessian matrix in Equation (10) can also be expressed in the form **η**′**H**^−1^**τ**. With these simplifications, the computation of the variance ratio *ω_i_* for the *i*th SNP is less computationally intensive via the Newton-Raphson method. We estimate the variance ratio *ω_i_* for the *i*th SNP by equating the gradient functions to be zero using the Newton-Raphson technique. With our simplifications,

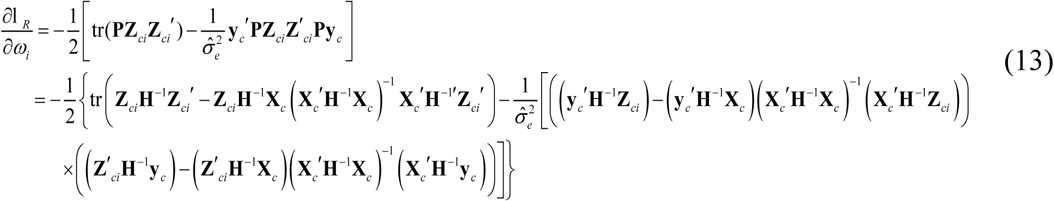

which is in the form **η**′**H**^−1^**τ**, and therefore its computation is so fast. We can also express the second derivative expression in Equation (10) in the form **η**′**H**^−1^**τ**:

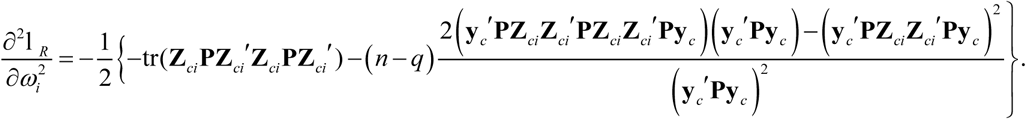

With these simplifications, the Newton-Raphson will converge smoothly to the estimate value of variance ratio *ω_i_* for the *i*th SNP. The gradient function and the Hessian matrix are in a simplified form. Therefore, the computations in each simulation run are fast. We have implemented this algorithm in R software.

**Empirical Bayes estimate of** *β_i_* : The joint distribution of **y** and *β_i_* is a multivariate normal distribution

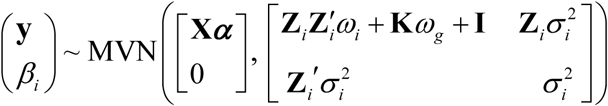

The conditional distribution of *β_i_* given **y** is

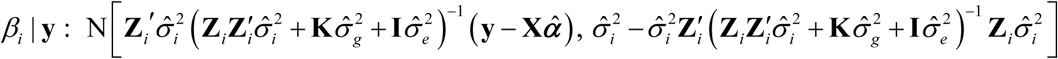

From a Bayesian analysis point of view, the conditional mean of *β_i_* given y is an empirical Bayes estimate of *β_i_*. Based on this framework, we obtain the Wald test statistic 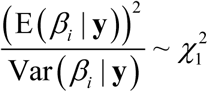 (Chi-square test with 1 degree of freedom) and using this distribution we obtain P-values for each marker effect. We test each marker effect at 1% level of significance. We do not perform multiple test correction because we intend to include markers that pass this initial test in a multi-locus model.

### Detection of true QTNs in multi-locus model

If the number of markers passing the 1% level of significance test is more than *n*, we invoke the LARS algorithm [32] to select the *n* –1 variables that are most likely associated with the quantitative trait of interest. LARS is a flexible method for variable selection which is conducted in lars package (http://cran.r-project.org/web/packages/lars/) in R language. The *n* − 1 markers are then included in a multi-locus model. Note that if the number of markers passing the initial test is less than *n*, we skip the LARS step and proceed to include all the selected markers in a multi-locus model. We compared various multi-locus methods: SCAD [34], adaptive Lasso [35] and EM-Empirical Bayes [36]. EM-Empirical Bayes [36] has the highest statistical power and accuracy of the estimated marker effects (Tables S8, S9, and S10). EM-Empirical Bayes is a random model method given as,

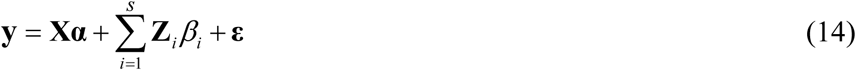

where **y**, **X** and **α** are the same as in model in Equation (1), *s* is the number of potentially associated markers selected from the first step in FASTmrMLM, **Z**_*i*_ and *β_i_* are *n*×1 incident vector and the random effect of the *i*th SNP, respectively. The polygenic variance is not included in the model because the model included all the potentially associated QTNs. We assume a normal prior for *β_i_*, 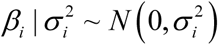 and a scaled *χ*^2^ prior for 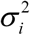, 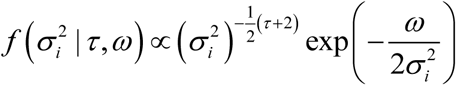 and we set (*τ*, *ω*) = (0,0), which is Jeffrey’s prior 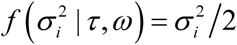 [36]. The procedure for parameter estimation in EM-Empirical Bayes is as follows:

a) Initial step: We set initial values as,

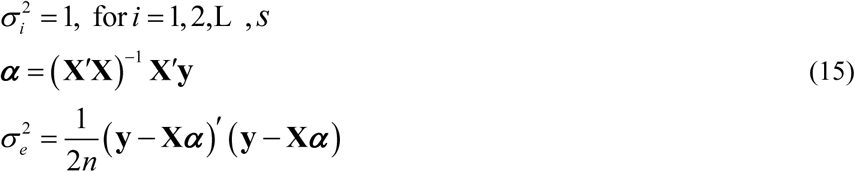
b) E-step: QTN effect can be predicted by

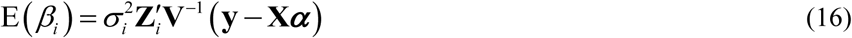

where 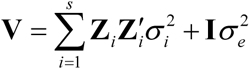.

c) M-step: To update parameters 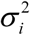, **α** and 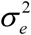

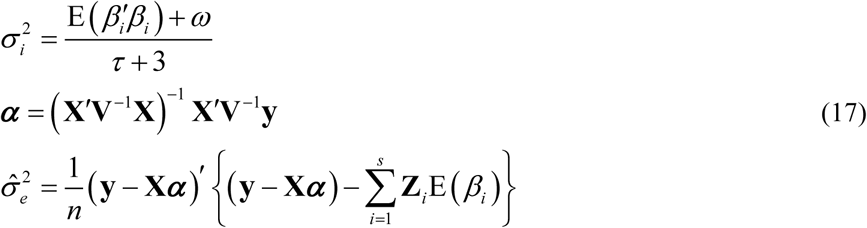

where 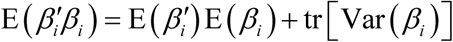, 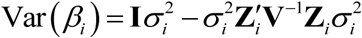 and (*τ*, *ω*) = (0, 0).

We repeat E-step and M-step until convergence is satisfied. We select all SNPs with a score LOD ≥ 3 (log of odds) and regard them as significant. We term our algorithm as a fast multilocus random-SNP-effect mixed linear model (FASTmrMLM).

### mrMLM

mrMLM is a two-stage method which tests each random marker effects in the first stage before estimating the significant putative QTNs in a multi-locus model [16]. mrMLM and FASTmrMLM methods were implemented by the R software mrMLM, which is downloaded from http://cran.r-project.org/web/packages/mrMLM/index.html. In mrMLM, we select all markers with a score LOD ≥ 3 and regard them as significant.

### GEMMA and EMMA

EMMA is a single-locus testing method for GWAS [6] and considers marker effects as fixed. The R codes for EMMA can be downloaded from http://mouse.cs.ucla.edu/emma/. GEMMA is a fast version of EMMA [14]. GEMMA can be run in Linux using source codes obtained from www.xzlab.org/software.html. For EMMA and GEMMA we select significant markers based on Bonferroni correction for multiple tests by setting a threshold for P-value at 0.05/*m*, where *m* is the number of markers.

### FarmCPU

FarmCPU was proposed by Liu et al. [28]. In this method, the fixed effect model and random effect model are used iteratively until a stage of convergence is reached. FarmCPU software package source codes are available at http://www.ZZLab.net/FarmCPU. We select significant markers based on Bonferroni correction for multiple tests by setting a threshold for P-value at 0.05/*m*, where *m* is the number of markers.

### SCAD

SCAD [34] is a shrinkage method that performs variables selection using concave penalties. SCAD can be run in R using the ncvreg package in R language downloaded from http://cran.r-project.org/web/packages/ncvreg/. Here we select all markers with a score LOD ≥ 3 and regard them as significant.

### Adaptive Lasso

Adaptive Lasso [35] is a variable selection method that uses data-dependent weights for *L*_1_ - penalizing coefficients in the penalty by choosing the inverse of ordinary least-square estimates for the weights. Adaptive Lasso can be run in R using the parcor package in R language downloaded from http://cran.r-project.org/web/packages/parcor/. Here we select all markers with a score LOD ≥ 3 and regard them as significant.

### Monte Carlo simulation experiments

We carried out three Monte Carlo simulation experiments to validate FASTmrMLM. In all these simulation studies, we sampled the genetic marker values from 216,130 SNPs markers in Atwell et al. [31] (http://www.arabidopsis.org/), and we simulated all the phenotypes values for quantitative traits. We used the same sample size of 199 diverse inbred lines like that in Atwell et al. [31] dataset. We randomly sampled 2000 SNPs from each of the five chromosomes; in total, we obtained 10000 SNP genotypes. For the selected SNPs, we used the same positions and genotypes as those in Wang et al. [16].

The first experiment tested FASTmrMLM method on a model with no polygenic variances simulated. From the sampled SNPs, we set six QTN with effect sizes and positions as shown in Tables S8 to S10. We placed these QTNs on the SNPs with allelic frequencies of 0.30. The heritabilities for these QTNs were 0.10, 0.05, 0.05, 0.15, 0.05 and 0.05, respectively. In this case, the phenotypes were simulated from the model: 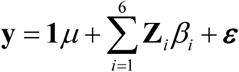, where ***ε*** : MVN(0,10×**I**). We simulated an overall mean of 10. In this simulation experiment, we carried out 1000 runs for each of the methods considered, i.e., FASTmrMLM, EMMA, GEMMA, FarmCPU, and mrMLM. For each simulated QTN, we counted the runs in which we obtained the QTN LOD ≥ 3 for FASTmrMLM and mrMLM and P-values less than 0.05*/m* (Bonferroni correction for multiple tests), where *m* is the total number of SNPs for GEMMA, EMMA, and FarmCPU. We considered QTNs within 1kb of the simulated QTN as true QTN. We counted the number of runs in which we deemed to have obtained each QTN. The proportion of such runs relative to the total number of runs (1000) represented the statistical power of this QTN. We calculated false positive rate (FPR) as the ratio of the number of false non-zero effects relative to the total number of zero effects considered in the full model. We measured the bias of each QTN effect estimate by calculating the mean squared error (MSE). We also carried out a paired t-test for the differences of statistical power or MSE between FASTmrMLM and other methods.

The second simulation experiment tested FASTmrMLM method on a model with an additive polygenic (small effect genes) background. We simulated polygenic effect from a multivariate normal distribution, 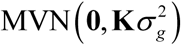 where 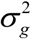 is the polygenic variance and **K** is the kinship matrix. We used 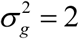 hence 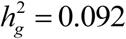. The QTN size (*h*^2^), average, residual variance, and other values were the same as those in the first simulation study, i.e., the phenotypes were simulated from the model given as: 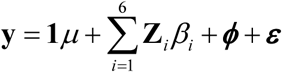, where ***ϕ*** ~ MVN(**0**, 2 × **K**) and ***ε*** ~ MVN(**0**,10 × **I**).

The third simulation experiment examined FASTmrMLM on a model with an epistatic background. We simulated three epistatic QTN each with 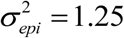 and 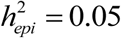. The details for the three epistatic QTN were described in Wang et al. [16]. The QTN size (*h*^2^), average, residual variance, and other values were the same as those in the first simulation experiment. The phenotypes were simulated from the model: 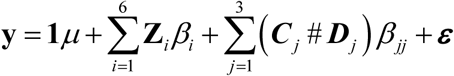, where, ***ε*** : MVN(0,10×**I**), *β_jj_* is the epistatic effect and (***C**_j_* # ***D**_j_*) is its incidence coefficient.

### The *Arabidopsis thaliana* data

We re-analyzed six flowering time-related traits in *Arabidopsis* [31] by GEMMA, FASTmrMLM, EMMA, FarmCPU and mrMLM to validate our new method. These traits are days to flowering under long days (LD), days to flowering under long days with vernalization (LDV), days to flowering under short days (SD), days to flowering under LD with no vernalization (0W), days to flowering under long days with 2 weeks vernalized (2W), and days to flowering under long days with 4 weeks vernalized (4W). We downloaded these datasets from http://www.arabidopsis.usc.edu/. In the real data analyses, we determined the significantly associated SNPs by the critical threshold of LOD ≥ 3 for FASTmrMLM and mrMLM, and with the P-values less than 0.05*/m* for GEMMA, EMMA, and FarmCPU. We mined candidate genes for the trait under study within 30 kb of the significantly relevant SNPs.

## Additional file

**Table S1.** Comparison of the Statistical power in the detection of QTN in three simulation experiments using five GWAS approaches

**Table S2.** Paired t-test for the differences of statistical power and mean squared error (MSE) between FASTmrMLM (new) and other methods

**Table S3.** Comparison of the Mean squared error (MSE) for QTN effects across 1000 replicates in three simulation experiments using FASTmrMLM, mrMLM, FarmCPU, and GEMMA/EMMA GWAS approaches

**Table S4.** Comparison of the Mean for QTN effects across 1000 replicates in three simulation experiments using FASTmrMLM, mrMLM, FarmCPU and GEMMA/EMMA GWAS approaches

**Table S5.** Goodness of fit (AIC, BIC) for SNPs detected by FASTmrMLM, mrMLM, FarmCPU, and GEMMA/EMMA where a lower value indicates a better fit **Table S6.** GWAS for six flowering time traits in Arabidopsis thaliana using FASTmrMLM, mrMLM, FarmCPU, and GEMMA methods

**Table S7.** New Genes detected only by FASTmrMLM in the GWAS of six flowering time traits in Arabidopsis thaliana

**Table S8.** Comparison of EM-Empirical Bayes, Adaptive Lasso, and SCAD with no polygenic variance simulated

**Table S9.** Comparison of EM-Empirical Bayes, Adaptive Lasso, and SCAD in the second simulation experiment with an additive polygenic background (explaining 0.092 of the phenotypic variance)

**Table S10.** Comparison of EM-Empirical Bayes, Adaptive Lasso, and SCAD in the third simulation experiment with three epistatic QTNs each explaining 0.05 of the phenotypic variance

## Abbreviations

GWAS: genome-wide association study
SNP: single nucleotide polymorphisms
QTN: quantitative trait nucleotide
QTL: quantitative trait locus
EM: expectation and maximization
EMMA: efficient mixed-model association
GEMMA: Genome-wide efficient mixed-model association
FarmCPU: fixed and random model circulating probability unification
mrMLM: multi-locus random-SNP-effect mixed linear model
FASTmrMLM: a fast mrMLM multilocus mixed linear model.
ISIS EM-BLASSO: Iterative modified-Sure Independence Screening EM-Bayesian LASSO
FASTmrEMMA: fast multi-locus random-SNP-effect EMMA.

## Authors’ contributions

Y.-M.Z. conceived and supervised the study, and revised the manuscript. C. L.T. performed the experiments and analysed the data, and wrote the draft. All the authors read, revised and approved the final manuscript.

## Acknowledgements

This work was supported by the National Natural Science Foundation of China (31571268), and Huazhong Agricultural University Scientific & Technological Self-innovation Foundation (Program No. 2014RC020).

## Availability of data and materials

A single source file for all simulated datasets has been included as Additional files.

